# Role of misfolded tau in the onset and progression of brain toxicity after trauma

**DOI:** 10.1101/2020.06.18.159301

**Authors:** Elisa R. Zanier, Maria Monica Barzago, Gloria Vegliante, Margherita Romeo, Ilaria Bertani, Carmina Natale, Luca Colnaghi, Laura Colombo, Luca Russo, Edoardo Micotti, Luana Fioriti, Roberto Chiesa, Luisa Diomede

**Affiliations:** Department of Neuroscience, Istituto di Ricerche Farmacologiche Mario Negri IRCCS, Milan, Italy; Department of Molecular Biochemistry and Pharmacology, Istituto di Ricerche Farmacologiche Mario Negri IRCCS, Milan, Italy

**Keywords:** tau, tauopathy, prion, traumatic brain injury, *Caenorhabditis elegans*

## Abstract

Traumatic brain injury (TBI) is associated with widespread tau pathology in about one third of patients. We previously found that TBI induces a transmissible tau pathology (tau^TBI^), with late cognitive decline and synaptic dysfunction. To understand whether tau^TBI^ is a marker of ongoing neurodegeneration or a driver of functional decline, we employed *C. elegans*. Brain homogenates from chronic TBI mice, or from mice in which tau^TBI^ had been transmitted by intracerebral inoculation, impaired *C. elegans* motility and neuromuscular synaptic transmission. Brain homogenates from tau P301L transgenic mice, or pre-aggregated recombinant tau, induced a similar toxic response. Protease digestion or pre-incubation of homogenates with anti-tau antibodies abolished toxicity, and TBI brain homogenates from tau knock-out mice had no toxic effect. These results support a vital role of abnormal tau species in chronic neurodegeneration after TBI and set the groundwork for the development of a *C. elegans*-based platform for screening anti-tau compounds.

## INTRODUCTION

Traumatic brain injury (TBI) is a leading cause of death and disability in young adults worldwide^1,2^. There is a growing acceptance that TBI is an acutely initiated event that results in long-lasting physical, and mental disability. In a sizeable minority TBI may also be a progressive disease, with worsening neuroimaging findings and neurology over months and years, and/or an increased risk of late dementia^3,4^. In the last decade, studies have led to the characterization of the neuropathology in individuals exposed to TBI of varying severity, and across injury survivors, from hours to many decades. These studies have given a complex picture of pathologies, with abnormalities in tau, amyloid beta (Aβ), neuronal loss, axonal degeneration, neuroinflammation and blood-brain barrier disruption – all changes common to wider neurodegenerative diseases^5–8^. The development of hyperphosphorylated tau (p-tau) is a hallmark of Alzheimer’s disease, correlating to outcome and late symptomatology^9^, and may be a crucial aspect linking TBI to late dementia. The recent addition of the positron emission tomography ligand flor-taucipir/11C-PBB3 to map tau pathology has provided information about the temporal and spatial characteristics of tau accumulation in the living brain, showing variable increases compared to healthy volunteers years after single or repetitive TBI^10–12^. However, in the clinical setting it is difficult to disentangle the precise significance of p-tau accumulation from other possible contributors to post-TBI neurodegeneration and/or dementia.

An important recent observation is that experimental TBI in wild-type (WT) mice induces the emergence and accumulation of a p-tau reminiscent of post-TBI tau pathology in humans^13,14^. Studies from our group indicate that this TBI-triggered p-tau pathology (tau^TBI^) spreads from the site of injury to remote regions of the brain and can be transmitted to naïve recipient mice by intracerebral inoculation, inducing memory deficits and synaptic alterations^14^. With the observation that tau^TBI^ is aggregated and partially resistant to pronase digestion, these results suggest that tau prions are generated in TBI, providing a possible explanation for how a biomechanical insult might trigger self-sustained neurodegeneration. However, it still needs to be established whether tau^TBI^ has a causal role in driving toxicity, as a primary player in the transition from an acute event to progressive neurodegenerative pathology.

To this aim we turned to the invertebrate *Caenorhabditis elegans*, which is widely used to investigate pathogenic mechanisms underlying protein-misfolding diseases, including tauopathies^15–20^. We established an integrated approach using brain tissue homogenates from TBI mice and *C. elegans* which allowed us to test directly whether tau^TBI^ is the primary toxic culprit in post-TBI neurodegeneration. We found that *C. elegans* motility and neuromuscular synaptic transmission were impaired when the nematodes were administered brain homogenates from chronic but not acute TBI mice, or from mice in which tau^TBI^ had been transmitted by intracerebral inoculation. There is a therapeutic opportunity to intervene after TBI before significant neurodegeneration takes place, so defining the pathogenic potential of TBI-generated tau proteoforms may have important clinical implications.

## RESULTS

### Brain-injured tissue from chronic TBI mice impairs *C. elegans* motility

We previously found that a single severe TBI induced persistent and evolving tau pathology in WT mice, which progressively spread from the site of injury to ipsi and contralateral brain regions. P-tau deposits were seen adjacent to the site of injury three months post-TBI, and were widespread in the ipsi and contralateral hemispheres at 12 months. Tau^TBI^ could be transmitted to naïve mice by intracerebral inoculation in a prion-like manner, causing synaptic toxicity and memory deficits^14^. To study tau^TBI^ toxicity in a more tractable model we used the nematode *C. elegans*, which is susceptible to the toxicity of various amyloidogenic proteins^21–24^.

Synchronized nematodes were incubated at L4 larval stage with brain homogenates (0.3 μg total proteins per worm) from TBI or sham mice euthanized at different times; naïve mice were used as an additional control (**Fig. 1a**). Locomotor activity and pharyngeal function were scored after 7 days. There was a significant reduction in the frequency of body bends in worms exposed to homogenates from chronic (3 and 12 months) but not acute (24h or 72h) TBI mice (**Fig. 1b**). Locomotor impairment was induced by brain tissue ipsilateral (TBIipsi) but not contralateral (TBI_contra_) to the site of injury collected three months post-TBI, and by both TBI_ipsi_ and TBI_contra_ tissues at 12 months (**Fig. 1b**), correlating with the presence of p-tau pathology^14^. The decrease in body bends was first detectable 2 days after treatment, and persisted at 4 and 7 days (**Fig. 1c**), exacerbating the physiological age-dependent decline in nematode motility (**Supplementary Fig. 1a**). No effect on pharyngeal pumping was observed at any time (**Fig. 1d**), indicating a selective neuromuscular defect. There were no changes in body bend frequency or pharyngeal pumping in worms incubated with brain homogenates from naïve or sham mice, compared to untreated controls (**Supplementary Fig. 1**).

**Fig. 1.**
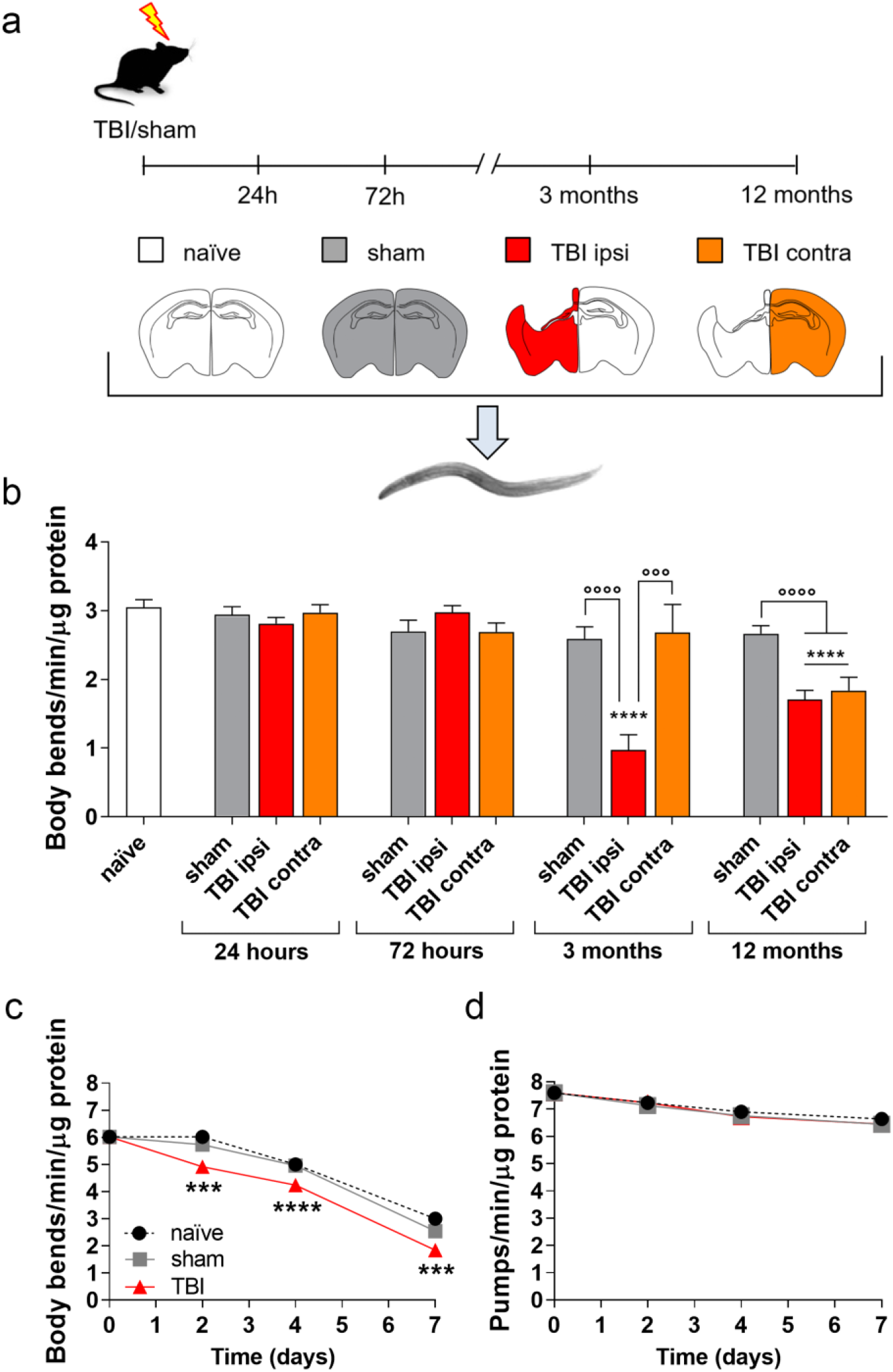
Brain homogenates from chronic but not acute TBI mice are toxic to *C. elegans*. **a** Pericontusional tissue homogenates were prepared from the ipsi (TBI ipsi) or contralateral (TBI contra) brain areas from TBI mice 24 h, 72 h, 3 months and 12 months post-injury. Brain homogenates from naïve and sham mice were used as controls. **b** Effects of the different brain homogenates on nematode locomotor activity 7 days after treatment. Data are mean + SEM of three independent experiments using brain homogenates from three different mice/group (50 worms/group). **** p<0.0001 *vs* naïve, °°°pBrain homogenates from chronic but not acute TBI mice are toxic to *C. elegans*.0.001 and °°°°p<0.0001, one-way ANOVA and Bonferroni’s *post hoc* test. Time-course of the effect on **c** locomotor and **d** pharyngeal activity of homogenates from 12-month TBI, sham or naïve mice (30 μg protein/100 worms/100 μL). Data are the mean ± SEM of three independent experiments with homogenates from two different mice/group (60 worms/group). ***p<0.001 and ****p<0.0001 *vs* sham at the corresponding time-point, one-way ANOVA and Bonferroni’s *post hoc* test

### Tau^TBI^ mediates toxicity in worms

We used different approaches to investigate whether the abnormal tau that accumulates in TBI mice at chronic stages was responsible for impairing the nematode locomotor function. First, we asked whether a deficit in motility like that induced by TBI homogenates was also seen in worms exposed to brain homogenates from P301L mice, which show robust tau pathology^25^. Worms were incubated with brain homogenates from nine-month-old P301L mice or age-matched Non-Tg controls, and body bend frequency and pharyngeal activity were scored after seven days. Like in worms treated with TBI tissue, the P301L homogenate significantly impaired body bend frequency but not pharyngeal activity (**Fig. 2**). The locomotor deficit worsened significantly on doubling the amount of P301L brain homogenate, consistent with a dose-dependent effect (**Supplementary Fig. 2**).

**Fig. 2.**
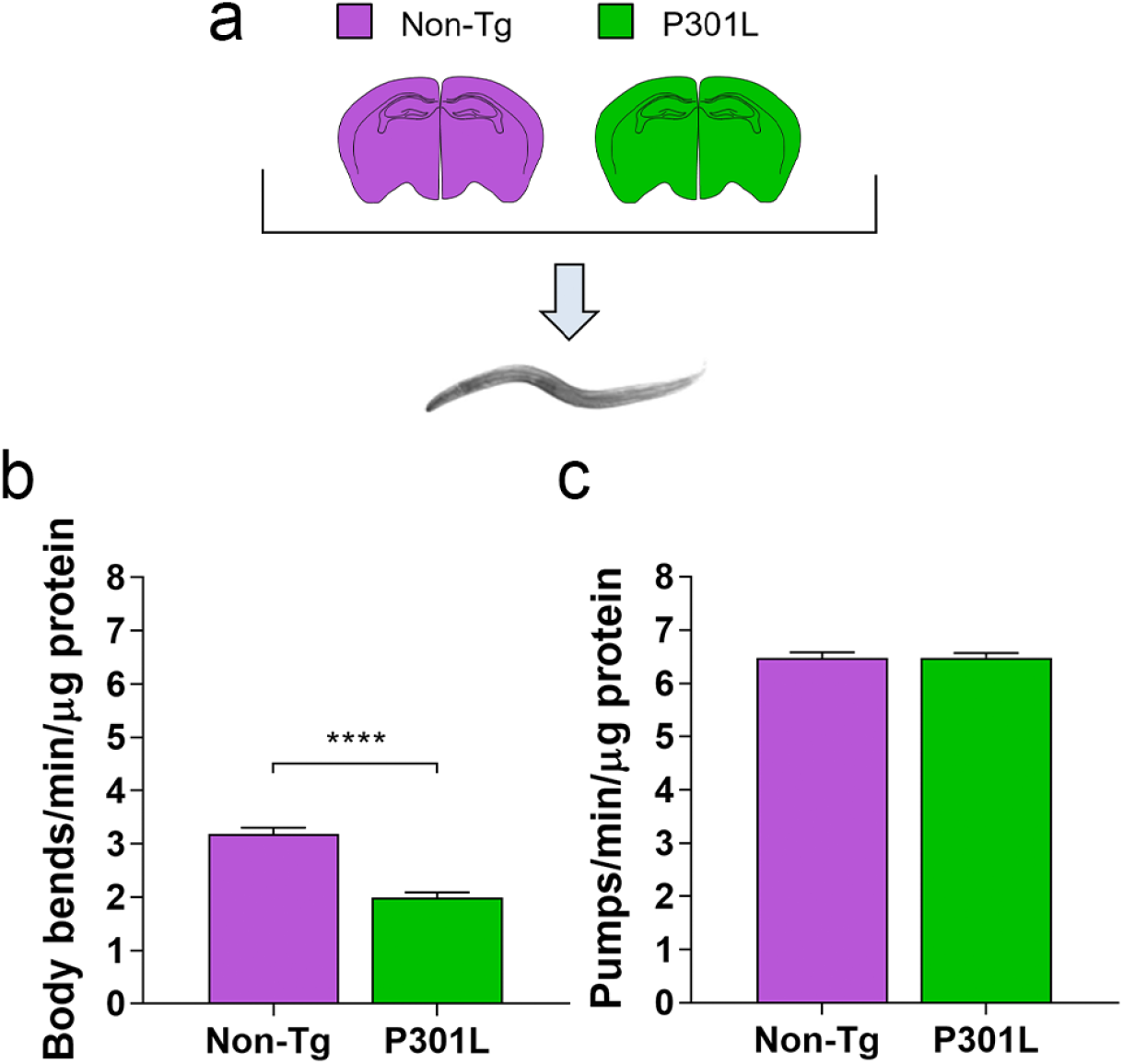
Brain homogenates from transgenic P301L mice are toxic to *C. elegans*. **a** Brains from Non-Tg or P301L mice were homogenized in 10 mM PBS, pH 7.4, and given to worms (30 μg protein/100 worms/100 μL). **b** Locomotor activity and (**c**) pharyngeal function of nematodes were rated 7 days after treatment. Data are the mean + SEM of three independent experiments with homogenates from three different mice (70 worms/group). **** p<0.0001 *vs* Non-Tg, Student’s t-test

Next, we asked whether the toxicity of TBI and P301L brains was abolished by treating them with high concentrations of PK, a broad-spectrum serine protease (**Fig. 3a, b**). Homogenates from TBIipsi (hereafter TBI) were used in all the subsequent experiments. Protein analysis by SDS-PAGE followed by Coomassie staining indicated complete protein digestion (**Fig. 3c, d**), consistent with tau^TBI^ being completely degraded by harsh protease digestion. PK completely abolished the ability of brain homogenates from P301L and TBI mice to induce the locomotor defect (**Fig. 3e, f** and **Supplementary videos**).

**Fig. 3.**
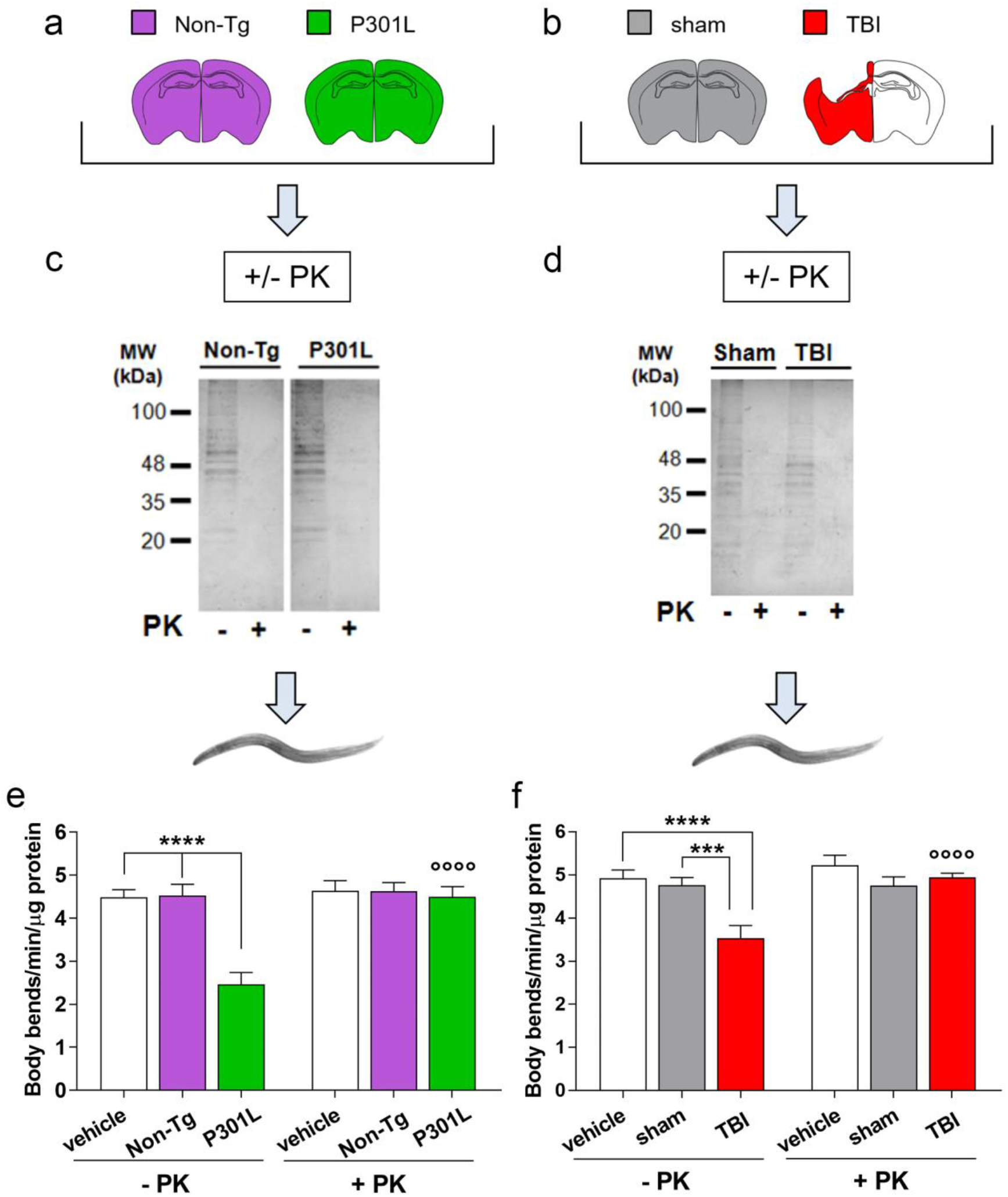
P301L and TBI brain homogenate toxicities are abolished by pretreatment with proteinase-K. Tissue homogenates (30 μg proteins) from **a** Non-Tg, P301L, **b** 12 months TBI and sham mice, were incubated for 1h at 37°C with 10 μg/mL proteinase-K (+PK) or the same volume of water (-PK). **c, d** Protein digestion was evaluated by SDS-PAGE analysis and staining with Comassie Blue (2 μg proteins/lane). **e, f** The same samples were given to *C. elegans* (30 μg proteins/100 worms/100 μl) and the locomotor activity was rated 7 days later. Control worms were treated with 100 μl of water ± 10 μg/mL PK (vehicle). Data are the mean + SEM (50 worms/group). **e** ****p<0.0001 and °°°°p<0.0001 *vs* P301L-PK, by one-way ANOVA and Bonferroni’s *post hoc* test. Interaction 0.0017, two-way ANOVA and Bonferroni’s *post hoc* test. **f** ***p<0.001, ****p<0.0001 and °°°° p<0.0001 *vs* TBI-PK, one-way ANOVA and Bonferroni’s *post hoc* test. Interaction <0.0001, by two-way ANOVA and Bonferroni’s *post hoc* test

We used several complementary approaches to directly assess the role of tau. First, to see whether anti-tau antibodies could block P301L and TBI homogenate toxicities, we treated the nematodes with P301L or TBI brain homogenates preincubated with or without the T46 antibody, which recognizes the C-terminal region of both human and mouse tau, or the SP70 antibody recognizing the N-terminal domain of human but not mouse tau (**Fig. 4**). Incubation with native but not heat-inactivated T46 antibody prevented the motility defect caused by the P301L and TBI brain homogenates (**Fig. 4c, e** and **Supplementary Fig. 3**). P301L toxicity was also abrogated by the human tau-specific SP70 antibody (**Fig. 4d**). These results point to direct involvement of tau in toxicity.

**Fig. 4.**
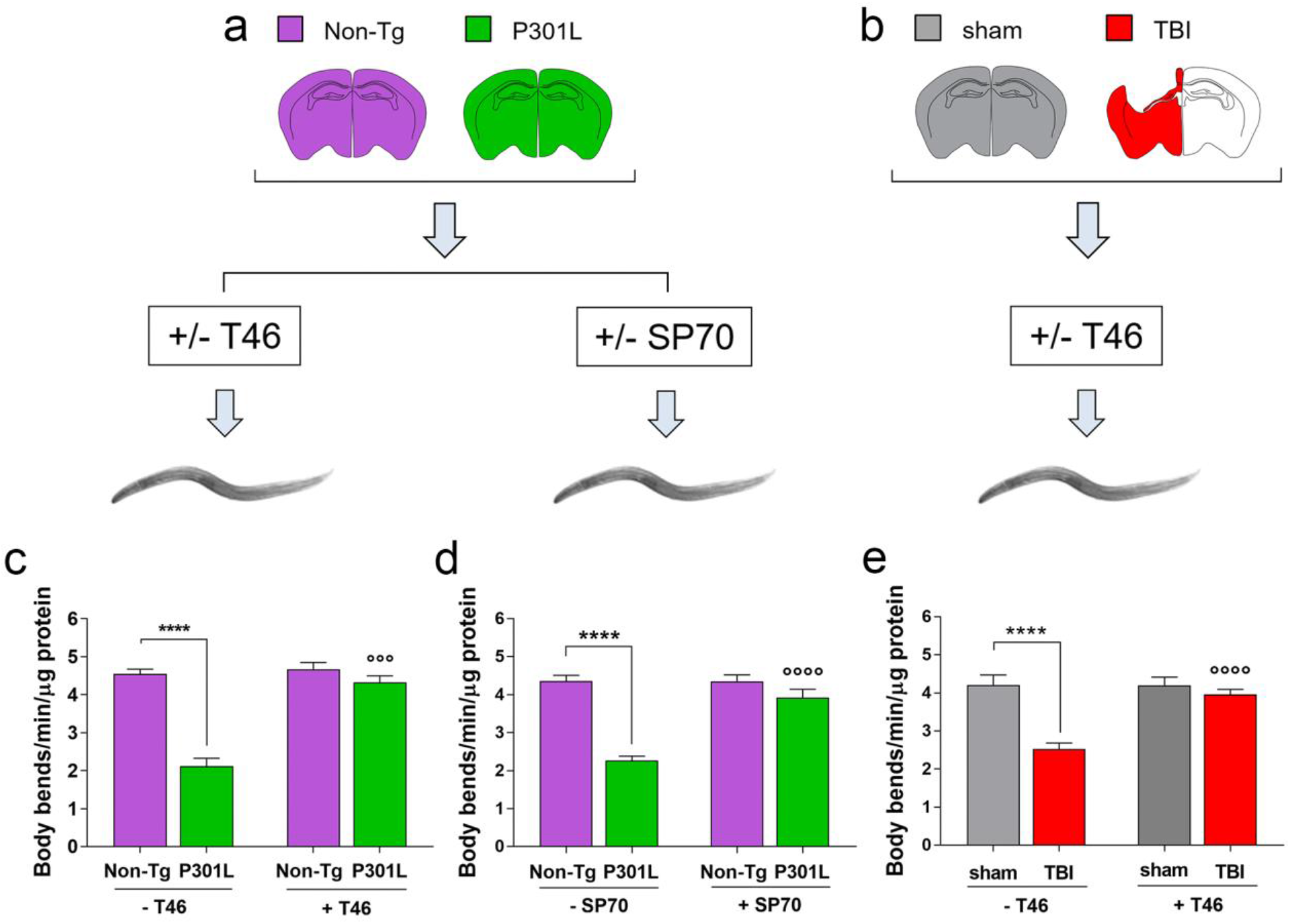
Anti-tau antibodies abolish P301L and TBI brain homogenate toxicities. Tissue homogenates (30 μg proteins) from **a** Non-Tg, P301L, **b** 12 months TBI and sham mice, were incubated for 30 min at room temperature with 10 ng/μL monoclonal anti-tau T46 antibody (+T46) or the same volume of water (-T46) or 5 ng/μL monoclonal anti-tau SP70 antibody (+SP70) or the same volume of water (-SP70). The homogenates (30 μg proteins/100 worms/100 μL) were then given to worms and locomotor activity was rated seven days later. Data are mean + SEM (50 worms/group). **c** and **d** ****p<0.0001 and °°°° p<0.0001 *vs* P301L-T46 or P301L-SP70, Interaction <0.0001, **e** **** p<0.0001 and °°°° p<0.0001 *vs* TBI-T46, Interaction=0.0008, according to two-way ANOVA and Bonferroni’s *post hoc* test

Next, we tested the toxicity of chronic TBI brain homogenates from tau KO mice. WT and tau KO mice were subjected to TBI or sham injury and euthanized after 4 months. There were no differences between WT and tau KO mice in TBI-induced sensorimotor deficits rated with the neuroscore and SNAP tests at 1 and 4 months (**Fig. 5a, b**) or in contusion volume assessed by T2w-MRI at 4 months (**Fig. 5c, d**). *C. elegans* treated with TBI homogenates from WT mice had significantly fewer body bends than those treated with WT sham homogenates, while homogenates from TBI tau KO mice had no effect (**Fig. 5e, f**).

**Fig. 5.**
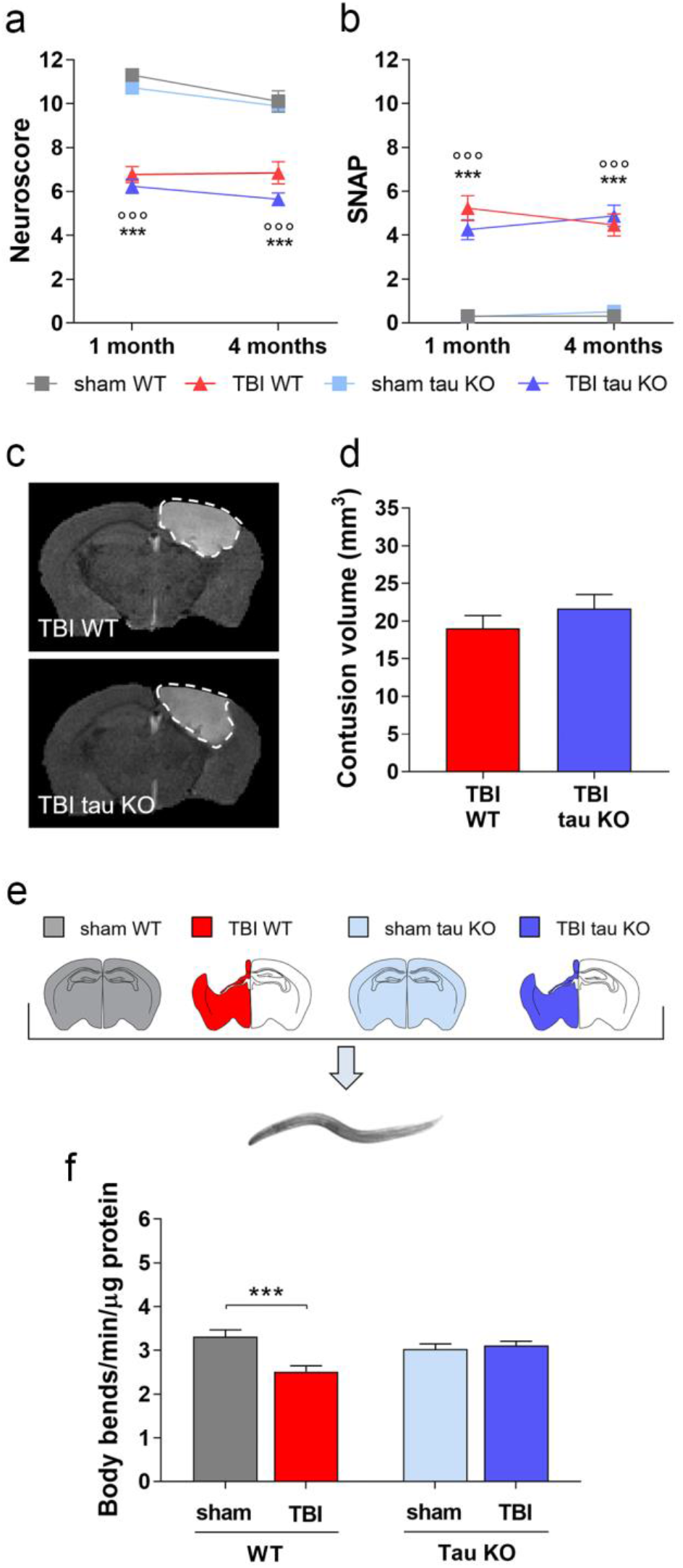
Brain homogenates from TBI tau knock-out mice are not toxic to *C. elegans*. Sensorimotor function of WT and tau knock-out (tau KO) mice was assessed by **a** neuroscore and **b** SNAP tests at 1 and 4 months after severe TBI or sham injury. Data are mean ± SEM, two way-ANOVA followed by Tukey’s *post hoc*. *** p<0.001 *vs* sham WT and °°° p<0.001 *vs* sham tau KO. **c**, **d** T2w-MRI images were used to evaluate contusion volume in TBI mice 4 months post-TBI. Data are mean + SEM, upaired t-test. **e** TBI homogenates of WT (TBI WT) or tau KO (TBI tau KO), sham WT or sham tau KO mice sacrificed 4 months post injury, were given to worms (30 μg protein/100 worms/100 μL). **f** The locomotor activity of nematodes was scored seven days later. Data are mean + SEM of data from three independent experiments with homogenates from three different mice (30 worms/group). *** p<0.001 *vs* sham, one-way ANOVA and Bonferroni’s *post hoc* test

Finally, we investigated whether exposing the nematodes to purified recombinant tau was sufficient to induce the deficit in body bends. Recombinant human 2N4R tau, either WT (Tau_WT_) or P301L (Tau_P301L_), was given to worms in the form of soluble monomers or insoluble aggregates obtained by incubating the proteins with heparin for 24h (**Fig. 6a**). There were no changes in motility in worms incubated with monomeric Tau_WT_ or Tau_P301L_, whereas the aggregated proteins significantly reduced body bends (**Fig. 6b**), but not pharyngeal activity (data not shown). The toxicity caused by aggregated tau was similar to that induced by P301L or TBI brain homogenates. These findings indicate that the toxic effect of tau on the *C. elegans* neuromuscular apparatus is related to its state of aggregation, independently from its phosphorylation, since the recombinant protein is not phosphorylated and further support a role for misfolded tau in the locomotor deficit caused by TBI brain homogenates.

**Fig. 6.**
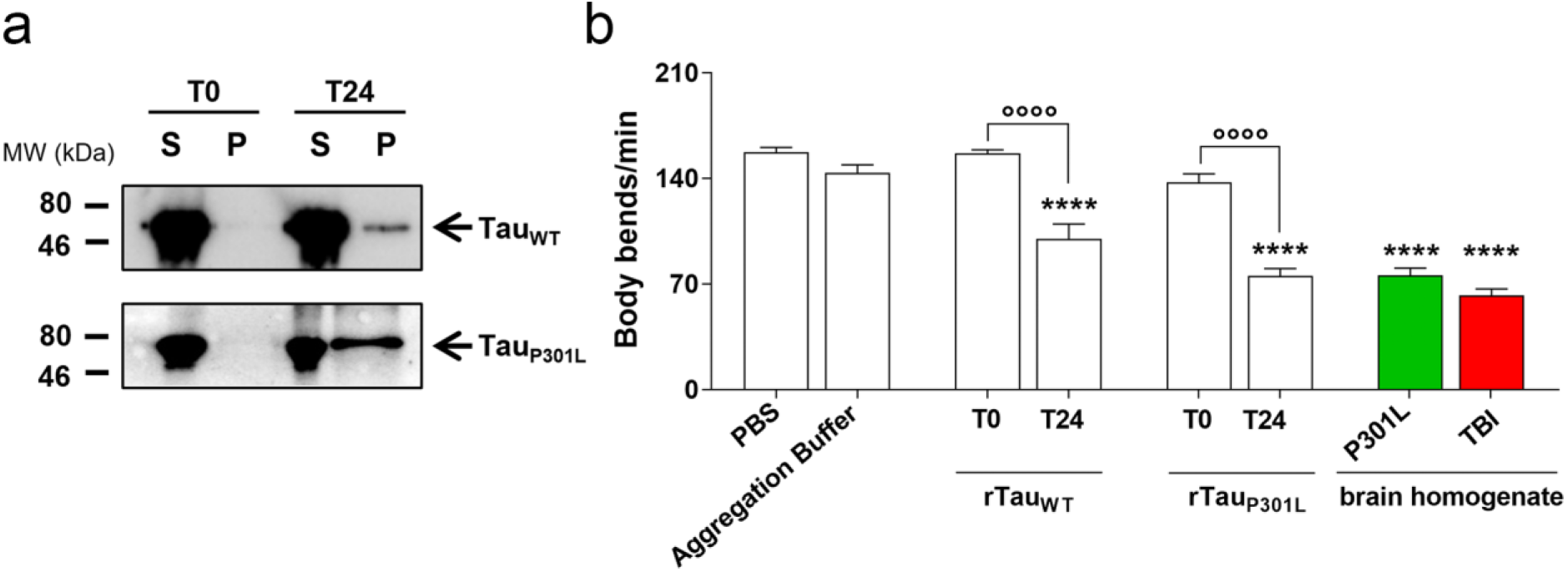
Aggregated recombinant tau is toxic to *C. elegans*. **a** A detergent insolubility test was done on TauWT and TauP301L immediately after addition of the co-aggregant heparin (T0) or after 24h of incubation at 37°C under shaking (T24). The supernatant (S) and pellet (P) were analyzed on 10% SDS-PAGE followed by Western blotting using the anti-total tau DAKO antibody. **b** An aliquot of the same proteins was diluted in 10 mM PBS, pH 7.4, and given to worms at the final concentration of 2 μg tau/100 worms/100 μL. *C. elegans* were treated in the same experimental conditions with 10 mM PBS, pH 7.4 (PBS) or co-aggregants diluted in 10 mM PBS, pH 7.4 (Aggregation buffer) as negative controls. Locomotor activity was rated 7 days after treatment. The effects of brain homogenates from P301L and 12 month TBI mice respectively (Figures 2b and 1b), are shown for comparison. Data are mean + SEM (50 worms/group). ****p<0.0001 *vs* PBS and °°°°p<0.0001, one-way ANOVA and Bonferroni’s *post hoc* test

### The locomotor impairment is due to neurotransmission failure

We investigated whether the locomotor dysfunction in nematodes treated with TBI or P301L brain homogenates could be ascribed to an impairment of synaptic transmission by analyzing the sensitivity to the paralyzing effect of aldicarb or levamisole. These pharmacological assays are commonly employed in *C. elegans* to depict the contribution of pre- and post-synaptic dysfunction to motility defects^26^. Aldicarb, an acetylcholinesterase inhibitor, causes the spastic paralysis of worms due to excessive acetylcholine build-up at the neuromuscular junction^26^. The speed of onset of paralysis depends on the acetylcholine released from the pre-synapse and the post-synapse competency in translating the cholinergic signal into a muscle contraction. Levamisole, a nicotinic acetylcholine receptor agonist, causes spastic paralysis by binding to acetylcholine receptors at post-synaptic sites on the neuromuscular junction^26^. Therefore, aldicarb gives information on the presence of both pre- and post-synaptic dysfunctions while levamisole indicates a post-synaptic defect.

The sensitivity to the paralyzing effect of aldicarb or levamisole was evaluated in nematodes treated with TBI, sham, P301L or Non-Tg homogenates or an equal volume of 10 mM PBS, pH 7.4 (vehicle).

Sensitivity to aldicarb was significantly greater in TBI-than in sham-treated nematodes and sensitivity to levamisole was reduced, indicating that both pre- and post-synaptic defects are responsible for the motility impairment (**Fig. 7a, b**). Results were similar in nematodes treated with brain homogenates from P301L mice (**Fig. 7c, d**).

**Fig. 7.**
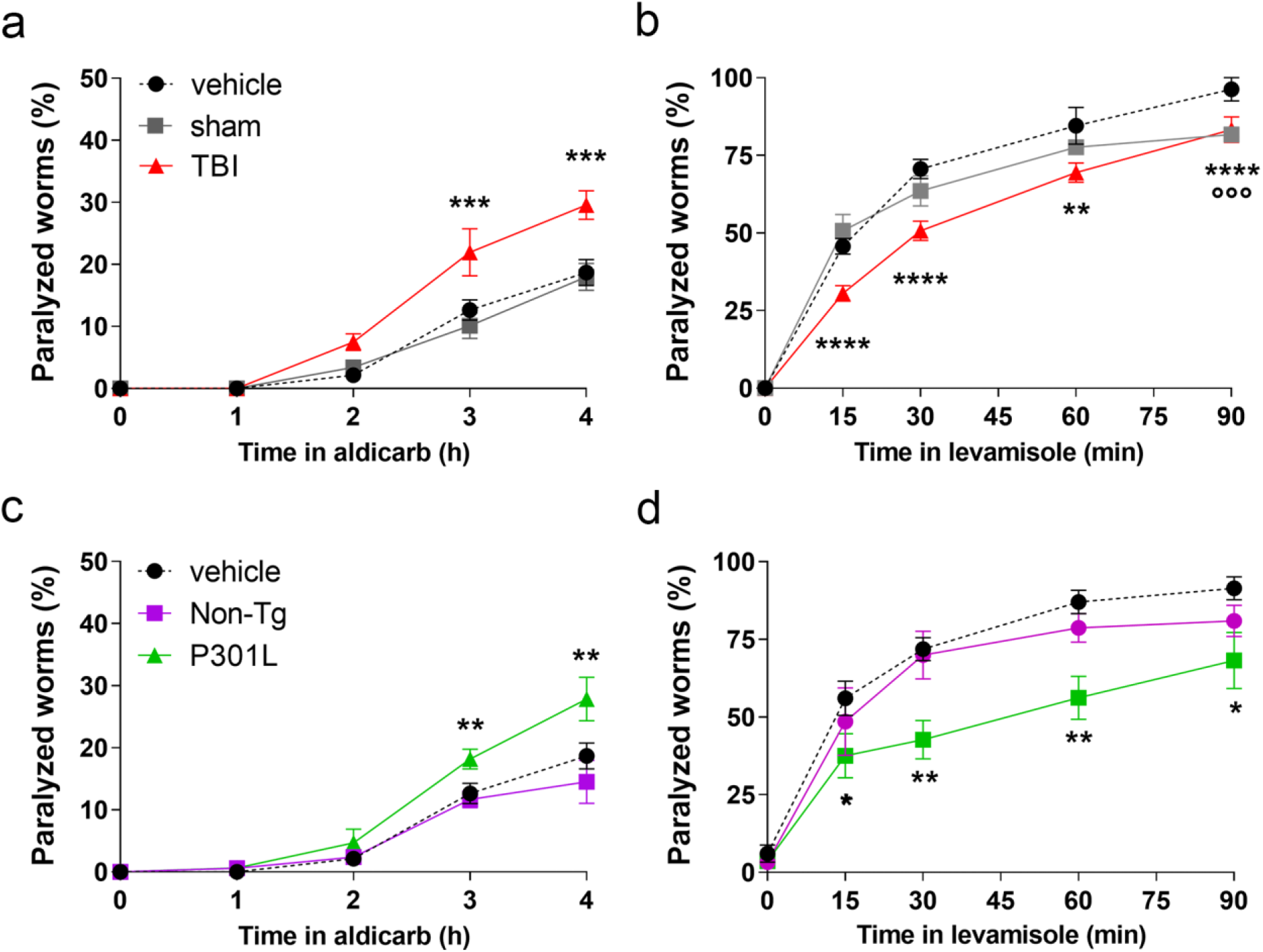
TBI and P301L brain homogenates induce pre- and post-synaptic defects. The sensitivity of worms to 0.5 mM aldicarb and 1 mM levamisole was evaluated 4 days after treatment with 10 mM PBS, pH 7.4 (vehicle) and brain homogenates (30 μg protein/100 worms/100 μL) from **a**, **b** 12 months post-TBI (TBI) and sham-injured (sham) mice or **c**, **d** Non-Tg and P301L mice. Data are mean ± SEM of percentages of paralyzed worms at different times after treatment with levamisole or aldicarb (100 worms/group). * p<0.05, **p<0.001, and ***p<0.001 *vs* Non-Tg or sham at the corresponding times according to one-way ANOVA and Bonferroni’s *post hoc* test

### Brain homogenates of mice replicating tau^TBI^ prions are toxic to *C. elegans*

We previously found that the hippocampus and thalamus are the two brain regions with the most prominent tau pathology in mice in which tau^TBI^ was transmitted by intracerebral inoculation ^14^. We asked whether brain homogenates from these two brain regions were toxic to nematodes. Homogenates were administered to worms and their locomotor activity was rated seven days later. There was a significant reduction in body bend frequency in worms treated with tau^TBI^-but not sham-inoculated homogenates (**Fig. 8**). The motility defect caused by brain homogenates from tau^TBI^-inoculated mice was comparable to that of worms treated with TBI homogenates (**Fig. 1**) indicating that the toxic properties of TBI-generated tau remained unaltered upon transmission and self-propagation in naïve mice.

**Fig. 8.**
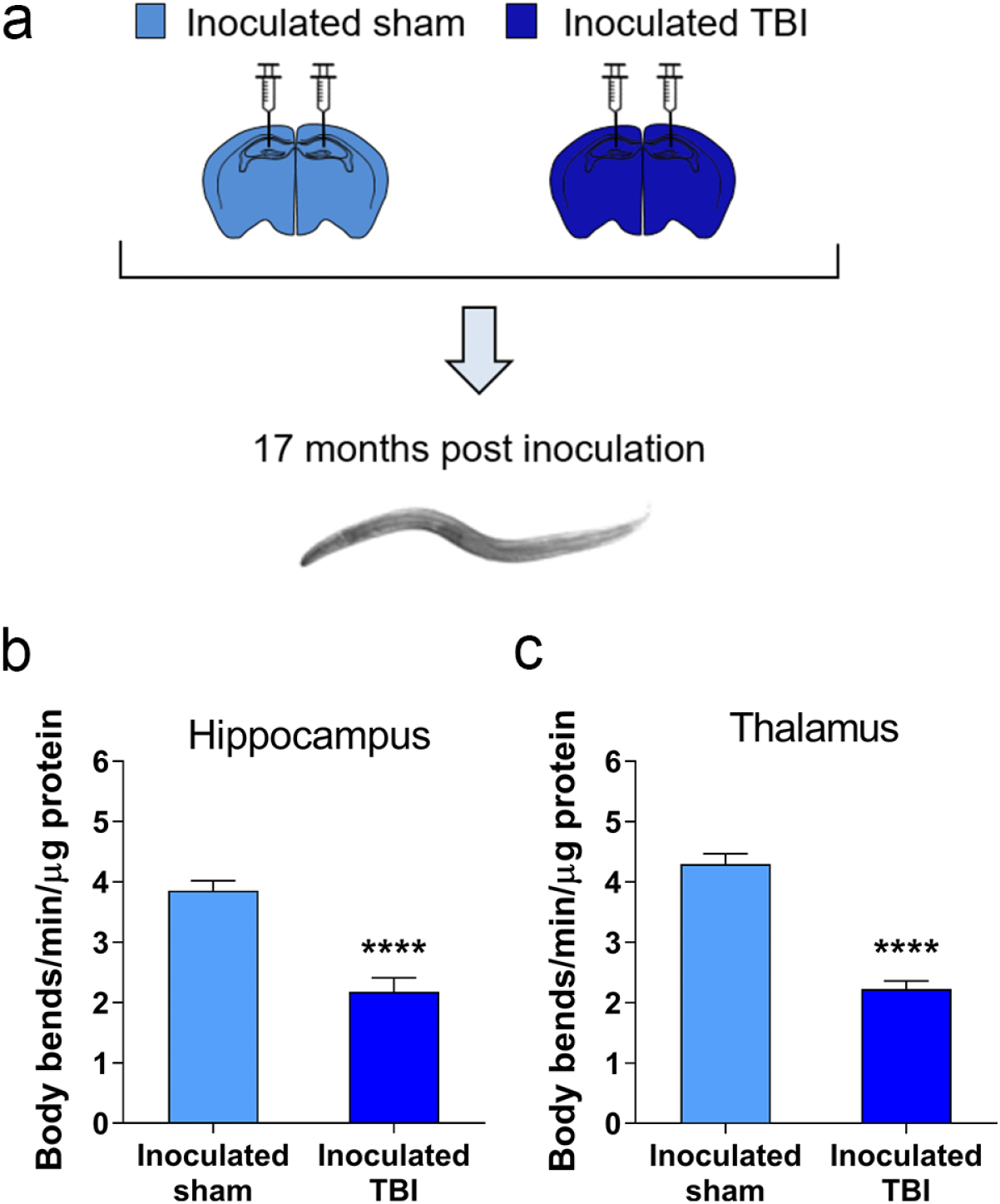
Brain homogenates from mice inoculated with contused brain tissue are toxic to *C. elegans*. **a** Homogenates of hippocampus or thalamus from mice inoculated with 12 months post-TBI (Inoculated TBI) or sham-injured (Inoculated sham) mice were given to *C. elegans* (30 μg protein/100 worms/100 μL). **b**, **c** Locomotor activity was rated 7 days after treatment. Data are mean + SEM from two independent experiments conducted with homogenates from three different mice (50 worms/group). **** p<0.0001 *vs* inoculated sham, according to one-way ANOVA and Bonferroni’s *post hoc* test

## DISCUSSION

This study shows that brain-injured tissue from chronic TBI mice impairs motility and synaptic transmission in *C. elegans*, and this is directly attributable to the abnormal form of tau that accumulates over time after TBI. These data indicate a pivotal role of abnormal tau conformers in chronic neurodegeneration after TBI.

*C. elegans* is widely used to investigate the *cell-autonomous* and *non-cell-autonomous* mechanisms underlying the pathogenesis of human diseases^20^. Its genes and pathways have a high degree of homology with those of vertebrates^15,27^ and its small size, short reproductive life cycle and lifespan allow to give large volumes of data in a short time^15^. In the context of protein misfolding neurodegenerative diseases and systemic amyloidosis, *C. elegans* has been used as a biosensor of toxic amyloidogenic proteins, as shown by our previous studies employing synthetic Aβ, HIV-1 matrix protein p17 peptides, amyloidogenic immunoglobulin light chains purified from patients’ biological fluids and recombinant gelsolin domains^21–24, 28^. This simple system has helped elucidate proteotoxic mechanisms and pave the way for novel therapeutic approaches^22,29^.

In the present study we exploited the biosensor properties of *C. elegans* to test the hypothesis that abnormal tau conformers play a causal role in driving toxicity in TBI. Chronic (3 and 12 months) but not acute (24 and 72 h) TBI tissue was toxic to nematodes, inducing a selective defect in locomotor activity. This indicates that toxicity is due to pathological changes occurring in the injured brain at chronic stages rather than to acute events, such as excitotoxicity, oxidative stress or acute inflammation. While at three months post-TBI only the ipsilateral brain homogenate was toxic, at 12 months both the ipsi and contralateral tissues induced the locomotor defect. This correlated with the progressive spread of tau pathology from the site of contusion to the ipsi- and contra-lateral brain hemispheres^13,14^.

This contention was further supported by the observation that brain homogenates from P301L mice, which develop genetically determined p-tau pathology with age, had similar deleterious effects on nematode neuromuscular function. The absence of toxicity in worms fed brain homogenates from sham or Non-Tg mice indicated the specificity of the effect and suggested that age-related brain changes other than p-tau accumulation had no role. The observation that the TBI and P301L brain homogenate toxicities were abolished by anti-tau antibodies, that brain homogenates from TBI tau KO mice were not toxic, and that recombinant tau was toxic to the worms, provided strong evidence of a direct role of tau. While in P301L mice proteinopathies other than tau have not been reported^25^, TBI favors misfolding of other proteins, accelerating Aβ and TDP-43 deposition^5,30^. Aβ administered to *C. elegans* is associated with transient pharyngeal impairment, but no motility deficits^21,28^, and in worms fed TBI tissue no pharyngeal impairment was observed. This confirms the primary role of tau in TBI-related toxicity and suggests that Aβ has a negligible effect if any.

*C. elegans* neuromuscular integrity and synaptic function deteriorate with age, contributing to the motility decline^31^. The defect in motility caused by TBI and P301L brain homogenates, however, was not due to hastened aging, since the lifespan of the nematodes was not affected (**Supplementary Fig. 4**). In aldicarb and levamisole sensitivity tests, the locomotor impairment was associated more with defective synaptic transmission, pointing to a synaptotoxic effect of tau. Transgenic *C. elegans* expressing human WT tau 2N4R in neurons accumulate insoluble forms of p-tau and, like worms fed TBI and P301L brain homogenates, have no impairment in pharyngeal function but develop locomotor and pre- and post-synaptic defects^32^. This suggests that the proteotoxic mechanisms of the neuromuscolar dysfunction induced by transgenically expressed and exogenously administered tau involve common molecular targets.

There is growing interest in the role of tau in post-TBI neurodegeneration, and the idea that targeting pathological tau may point to a therapeutic opportunity in TBI is growing^13,33–35^. The *C. elegans* model of tau^TBI^ toxicity described here could be employed to rapidly screen TBI conditions involving the generation of potentially toxic tau species in both experimental and human TBI, to investigate the mechanisms underpinning chronic neurodegeneration in TBI, and to develop a high-throughput experimental platform to screen for pharmacological agents that interfere with TBI-associated tau toxicity.

## METHODS

### Mice

Male C57BL/6J mice, referred to as WT mice, were purchased from Envigo (Holland). Male B6.129S4(Cg)-Mapt^tm1(EGFP)Klt^/J, referred to as tau knock-out (KO) mice, were from Jackson Laboratory (#029219, USA)^36^. Male JNPL3 mice expressing 0N4R human tau with the P301L mutation (P301L) were obtained from Taconic Biosciences (New York, USA) (Tau-Model 2508). Controls were non-transgenic (Non-Tg) male mice with the same mixed C57BL/6, DBA/2, SW genetic background as P301L mice. Mice were housed in a specific pathogen-free animal room at a constant temperature of 21 ± 1°C, humidity 60 ± 5%, with a 12h light/dark cycle, and *ad libitum* access to food and water.

Procedures involving animals and their care were conducted in conformity with the institutional guidelines at the Istituto di Ricerche Farmacologiche Mario Negri IRCCS in compliance with national (D.lgs 26/2014; Authorization n. 19/2008-A issued March 6, 2008 by Ministry of Health) and international laws and policies (EEC Council Directive 2010/63/UE; the NIH Guide for the Care and Use of Laboratory Animals, 2011 edition). They were reviewed and approved by the Mario Negri Institute Animal Care and Use Committee that includes ad hoc members for ethical issues, and by the Italian Ministry of Health (Decreto no. D/07/2013-B and 301/2017-PR). Animal facilities meet international standards and are regularly checked by a certified veterinarian who is responsible for health monitoring, animal welfare supervision, experimental protocols and review of procedures.

### Mouse model of TBI

The controlled cortical impact brain injury mouse model used in this study replicates both the mechanical forces and the main secondary injury processes observed in severe TBI patients with brain contusion and gives clinically-relevant behavioral and histopathological outcomes^14,37^. WT and tau KO mice were anesthetized by isoflurane inhalation (induction 3%; maintenance 1.5%) in an N2O/O2 (70%/30%) mixture and placed in a stereotaxic frame. Rectal temperature was maintained at 37°C. Mice were then subjected to craniectomy followed by induction of controlled cortical impact brain injury as previously described^38^. Briefly, the injury was induced using a 3-mm rigid impactor driven by a pneumatic piston rigidly mounted at an angle of 20° from the vertical plane and applied to the exposed dura mater, between bregma and lambda, over the left parietotemporal cortex (antero-posteriority: −2.5 mm, laterality: −2.5 mm), at an impactor velocity of 5 m/s and deformation depth 1 mm, resulting in a severe level of injury^39^. The craniotomy was then covered with a cranioplasty and the scalp sutured. Sham mice received identical anesthesia and surgery without brain injury. Mice were allocated to surgery by a list randomizer (http://www.random.org/list). All behavioral evaluations and magnetic resonance imaging (MRI) analysis were done blinded to injury status and genetic background.

### Sensorimotor function

TBI-induced sensorimotor deficits were assessed on WT and tau KO mice by the Neuroscore and Simple Neuroassessment of Asymmetric Impairment (SNAP) tests 4 months post-TBI. **Neuroscore.** Mice are scored from 4 (normal) to 0 (severely impaired) for each of the following indices: 1) forelimb function while walking on the grid and flexion function response when suspended by the tail; 2) hindlimb function while walking on the grid and extension function when suspended by the tail; and 3) resistance to lateral right and left pulsion. The maximum score per mouse is 12^38,40^. **SNAP.** The test evaluates eight neurological parameters including vision, proprioception, motor strength and posture. Score ranges from 0 (normal) to 5 (severely impaired) for each test. The scores from each of the eight tests are summed to give the total SNAP score. A neurologically intact animal would be expected to have a SNAP score of 0. The higher the score the more the asymmetric deficits^41^.

### Magnetic resonance imaging

Imaging studies were done four months after TBI. Images were acquired on a 7T Bruker Biospec (Ettlingen, Germany) running ParaVision 6.01, equipped with a quadrature cryogenic surface coil as transmitter and receiver. Mice were anesthetized (induction 3-4%, maintenance 1.5-2% in an air/O2 −70%/30% mixture) and body temperature was maintained at 37°C. Acquisitions: coronal, 2D, T2-weighted RARE sequences with FOV 1.5×1.5 cm, matrix 150×150, 37 slices 300 μm thick, repetition time TR 5500 ms, echo time E=66 ms, RARE factor of 8 and number of averages N=12. Contusion volume was computed using the ITK-SNAP software.

### Recombinant tau

Recombinant tau WT (Tau_WT_) and P301L (Tau_P301L_) were expressed in *E. coli* and purified by exchange chromatography followed by size-exclusion chromatography. The purification protocol was adapted from Rossi *et al*.^42^. Aggregation was induced by incubating the proteins with heparin (Mr 6000-20000, Selleckchem), in a tau:heparin ratio of 4:1 (w/w) in 10 mM phosphate buffered saline (PBS), pH 7.4, for 24h at 37°C with shacking. To assess aggregation, the samples were examined in a detergent insolubility assay adapted from Drisaldi et al.^43^ and the proteins in the supernatant and pellet were analyzed by Western blot using the anti-tau rabbit polyclonal antibody DAKO (Agilent Technologies Italy SpA, Milan, Italy) (1:10000 dilution). Anti-rabbit IgG peroxidase conjugate (1:20000, Sigma) was used as secondary antibody.

### Proteinase-K digestion

Brain homogenates (30 μg proteins) from the ipsi pericontusional area of 3 months post-TBI and sham-injured mice, Non-Tg and P301L were incubated for 1h at 37°C with 10 μg/mL proteinase-K (PK) or the same volume of water. After incubation, 2 μg of digested proteins were suspended in 4X SDS (sodium dodecyl sulfate) loading buffer, boiled for 5 min and analyzed in 10% SDS-PAGE, followed by Comassie Blue R250 staining for 30 min.

### *C. elegans* studies

Bristol N2 nematodes were obtained from the *Caenorhabditis elegans* Genetic Center (CGC, University of Minnesota, Minneapolis, MN, USA) and propagated at 20°C on solid Nematode Growth Medium (NGM) seeded with *E. coli* OP50 (CGC) for food. We used the bleaching technique to prepare age-synchronized animals^44^. *C. elegan*s at the first larval stage were then transferred to fresh NGM plates and grown at 20°C.

At L3-L4 larval stage nematodes were collected with M9 buffer, centrifuged, and washed twice with 10 mM PBS, pH 7.4, to eliminate bacteria. Worms were incubated for 2h at room temperature with orbital shaking, in the absence of *E. coli*, with homogenates from ipsi or contralateral (contra) areas from WT or tau KO TBI mice, the equivalent area of either WT or tau KO sham mice, or brain homogenates from Non-Tg and P301L mice (30-60 μg protein/100 worms/100 μL in 10 mM PBS, pH 7.4). Tau_WT_ or Tau_P301L_ (30 μg protein/100 worms/100 μL) were administered to worms in 10 mM PBS, pH 7.4. Worms incubated with brain homogenates of naïve mice (30 μg protein/100 worms/100 μL) or 10 mM PBS, pH 7.4 (100 worms/100 μL) were used as negative controls. Worms were then plated onto NGM plates seeded with OP50 *E. coli*, grown at 20°C and transferred every day to new NGM plates seeded with *E. coli* to avoid overlapping generations.

The locomotor activity and the pharyngeal function of nematodes were scored before (time 0) and 2, 4, and 7 days after the treatment^22,32^. To measure the locomotor activity, worms were picked up, transferred into a well of a 96-well ELISA plate containing 100 μL of ddH2O and their motility was evaluated by counting the number of left–right movements in 1 minute (body bends/min). The pharyngeal pumping rate was measured by counting the number of times the terminal bulb of the pharynx contracted in 1minute (pumps/min).

Brain homogenates (30 μg proteins) from the pericontusional area of 12 months post-TBI and sham-injured mice, Non-Tg and P301L were incubated for 1h at 37°C with 10 μg/mL PK or the same volume of water. The homogenates were then given to *C. elegans* (30 μg proteins/100 worms/100 μL) as described above. Control worms were treated with 100 μL of water ± 10 μg/mL PK. Worms were then plated onto NGM plates seeded with OP50 *E. coli*, grown at 20°C and transferred every day for six days to new NGM plates seeded with *E. coli*. Locomotor activity was rated on the seventh day after treatment.

In some experiments, N2 nematodes were fed for 2h with the different brain homogenates (30 μg proteins/100 worms/100 μL) previously incubated or not for 30 min at room temperature with 5-50 ng/μL anti-mouse tau monoclonal antibody T46 (Thermo Scientific) or anti-human tau monoclonal antibody SP70 (Rockland, Limerick, PA, USA). Antibodies alone were employed as controls. Worms were also treated in the same conditions with brain homogenates incubated for 30 min at room temperature with 10 ng/μL of T46 antibody inactivated by incubation at 100°C for 10 min (inactivated T46) or the corresponding volume of 10 mM PBS, pH 7.4. Worms were then plated onto NGM plates seeded with OP50 *E. coli*, grown at 20°C and transferred every day for six days to new NGM plates seeded with *E. coli*. Locomotor activity was rated on the seventh day. All behavioral evaluations were done blinded.

### Sensitivity to aldicarb and levamisole

The sensitivity of worms to aldicarb and levamisole was evaluated four days after treatment with brain homogenates. Nematodes (50 worms/plate) on NGM agar plates with OP50 *E. coli*, were treated with 0.5 mM aldicarb (Sigma Aldrich) or 1 mM levamisole (Sigma Aldrich). The worms were prodded on the nose after 15 min and 30-min intervals for the next 4h to determine when paralysis was complete^26^.

### Lifespan

For lifespan experiments, L4 worms were fed for 2h with the different brain homogenates, described above. Nematodes were then plated onto NGM plates seeded with OP50 *E. coli* and kept at 20°C. To avoid overlapping generations, live worms were transferred daily to fresh NGM plates seeded with OP50 *E. coli* without fluorodeoxyuridine until they were all dead. Nematodes were scored as dead when there was no touch-provoked movement. The first day of adulthood is day 1 in survival curves.

### Statistical analysis

The data were analyzed using GraphPad Prism 8.0 software (CA, USA) by Student’s t-test, one-way or two-way ANOVA and Bonferroni’s or Tukey’s post hoc test. The median survivals were determined using the same software. A p value < 0.05 was considered significant.

## Data availability

The data that support the findings of this study are available from the corresponding author, upon reasonable request.

## ACKNOWLEDGEMENTS

This work was supported by Fondazione Regionale per la Ricerca Biomedica (Care4NeuroRare CP_20/2018) to LD; the Alzheimer’s Association (AARG-17-532633 to ERZ and RC, AARG-17-505136 to LF; Fondazione Telethon (Telethon Career Award TCP15011) to LF. LC is the recipient of the BrightFocus award (A2019296F) and the Fondo di Beneficenza - Gruppo Intesa Sanpaolo. *C. elegans* and OP50 *E. coli* were provided by the GCG, which is funded by NIH Office Research Infrastructure Programs (P40 OD010440). The authors would like to thank Prof David K Menon for numerous useful discussions.

## CONTRIBUTIONS

Conceptualization: E.R.Z., R.C., and L.D.; Design and performance of experiments: M.M.B., G.V., M.R., and L.C.; Additional experimental work: I.B., C.N., L.COLO., L.R., and E.M.; Writing: E.R.Z., R.C., and L.D.; Data discussion and analysis: E.R.Z., R.C., L.D. and L.F.; Funding L.D., E.R.Z., R.C. and L.C.; Supervision: E.R.Z., R.C., and L.D.

## SUPPLEMENTARY MATERIAL

**Supplementary Fig. 1.**
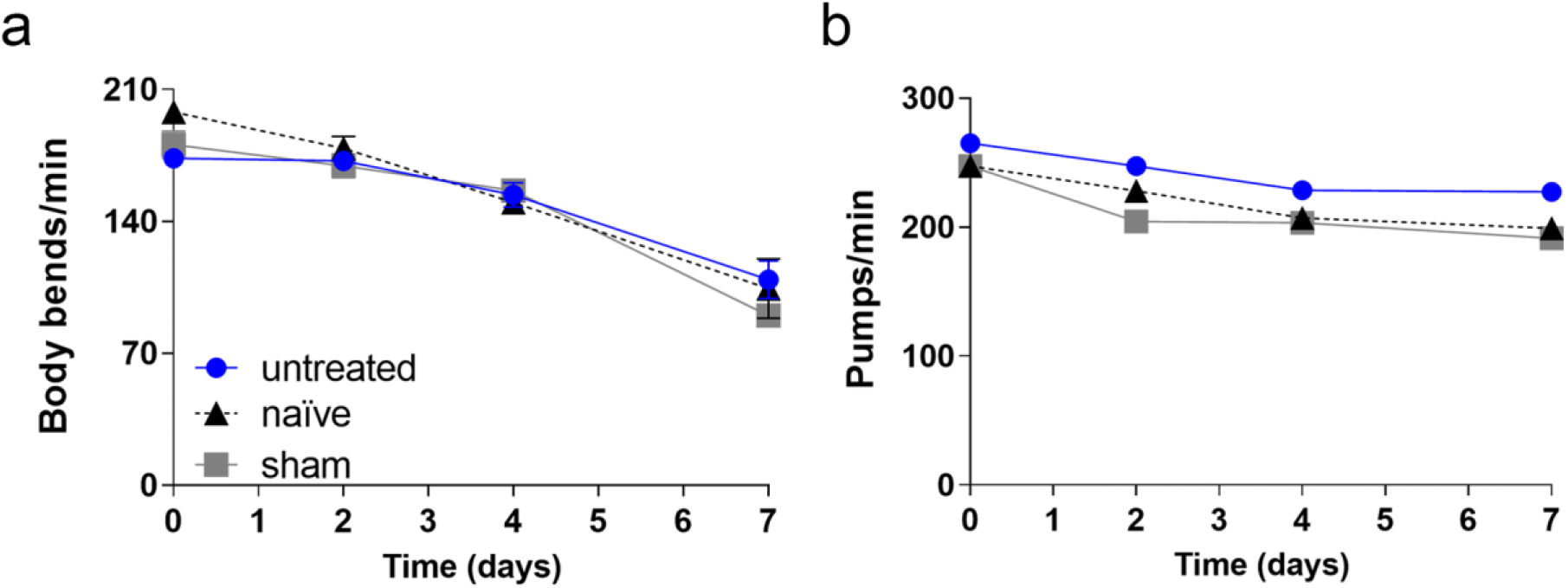
Sham brain homogenates did not modify the physiological age-dependent decline of the locomotor and pharyngeal activity of worms. N2 worms (100 worms/100 μL) were treated with 30 μg protein of brain homogenate from naïve or sham injured mice (sham). Untreated worms were used as control. The locomotor activity **a** and pharyngeal function **b** of nematodes were rated from day 1 to day 7 after treatment. Data are mean ± SEM of data from three independent experiments with homogenates from two different mice (30 worms/group)

**Supplementary Fig. 2.**
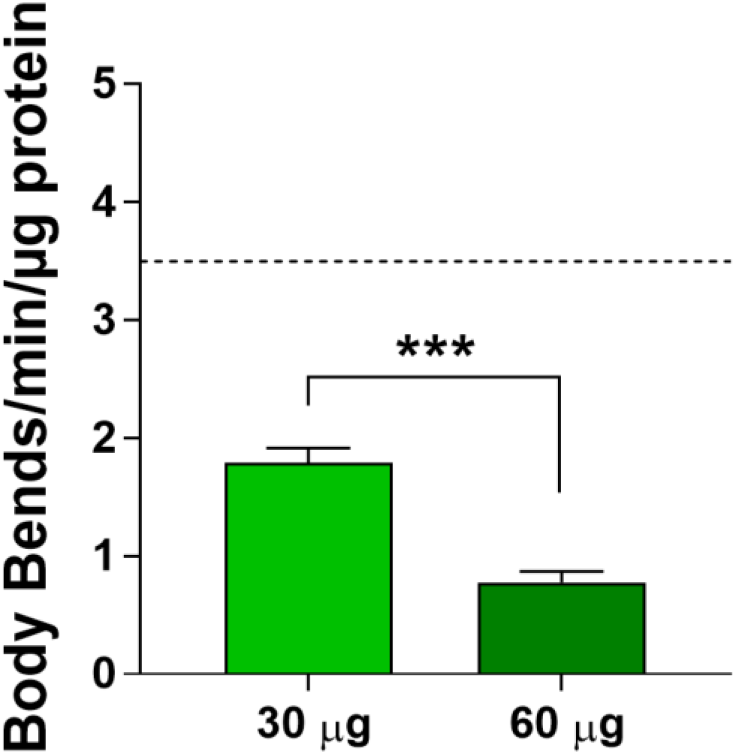
Dose-dependent effect of P301L brain homogenate on *C. elegans* locomotion. N2 worms (100 worms/100 μL) were treated with 30 or 60 μg protein of brain homogenate from Non-Tg (dotted line) or P301L transgenic mice (green column) and their locomotor activity was rated 7 days later. Data are mean ± SE (30 worms/group). ***p<0.001 according to one-way ANOVA and Bonferroni’s *post hoc* test

**Supplementary Fig. 3.**
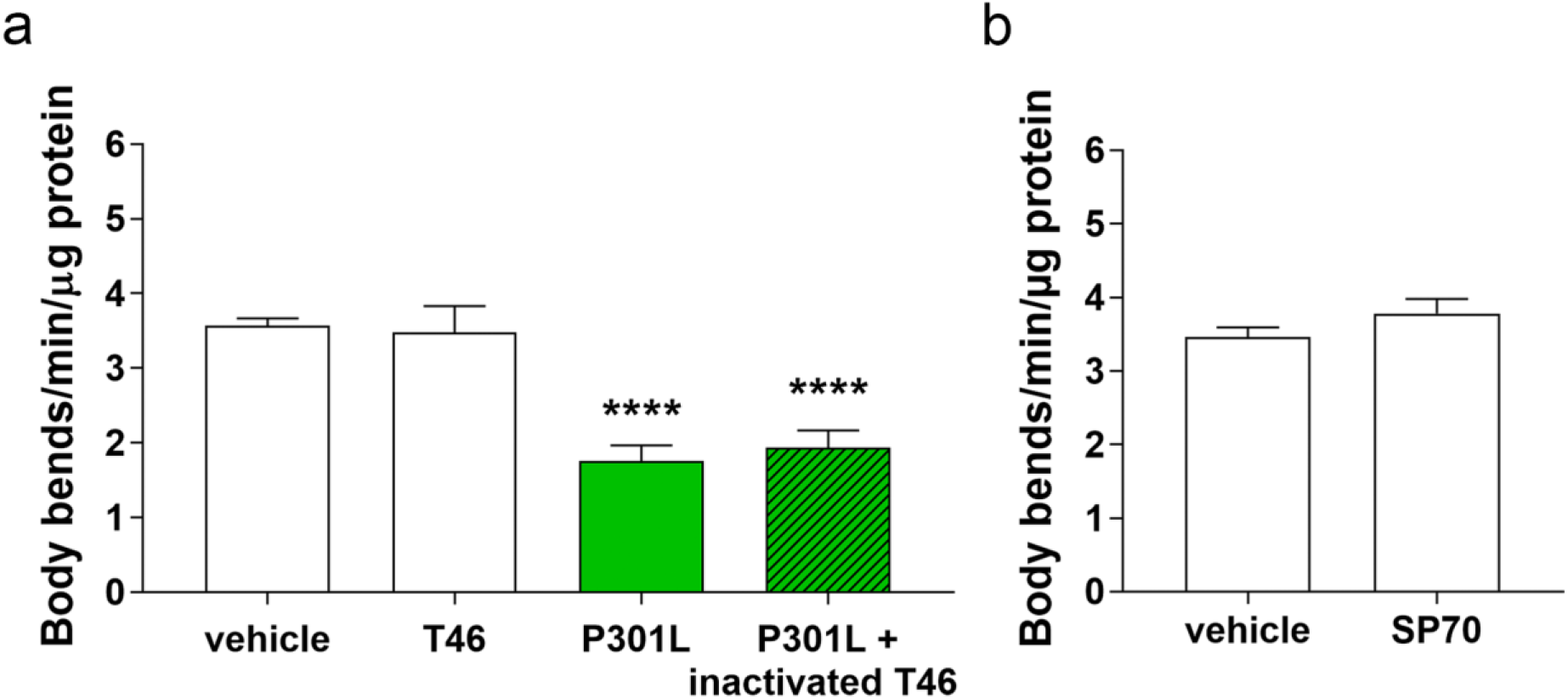
The anti-tau antibodies have no effect on *C. elegans* motility and heat inactivation abolished the antibody’s ability to rescue the P301L brain homogenate-induced locomotor defect. **a** The brain homogenate from P301L mice was administered to worms (30 μg proteins/100 worms/100 μl) before (P301L) or after 30 min pre-incubation for at room temperature with the monoclonal anti-tau T46 antibody (10 ng/μL) previously inactivated by incubation at 100°C for 10 min (P301L + inactivated T46). The effect of antibody alone was also examined by treating worms (100 worms/100 μL) in the same experimental conditions with **a** 10 ng/μL monoclonal anti-tau T46 antibody (T46) or **b** 5 ng/μL monoclonal anti-tau SP70 antibody (SP70). Control worms were treated with 10 mM PBS, pH 7.4 (vehicle). The locomotor activity of nematodes was rated seven days after the treatment. Data are mean ± SE. ****p<0.0001 *vs*. vehicle, one-way ANOVA and Bonferroni’s *post hoc* test

**Supplementary Fig. 4.**
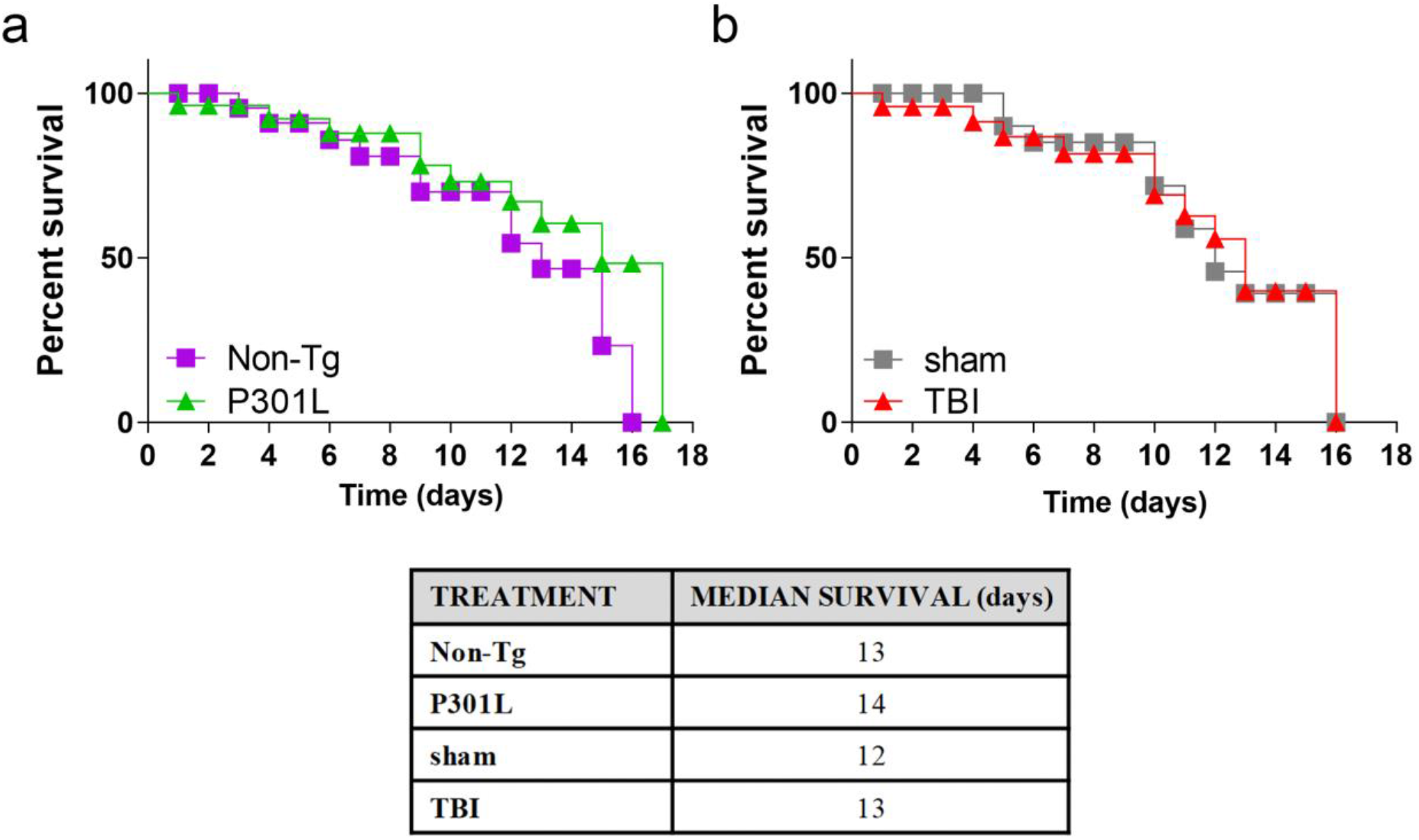
TBI and P301L brain homogenates do not affect *C. elegans* lifespan. **a** Brains from Non-Tg and P301L mice or **b** pericontusional tissue homogenates from the ipsilateral brain area of 12 months post-TBI mice (TBI) and brain homogenates from sham mice (sham), were homogenized in 10 mM PBS, pH 7.4, and given to worms (30 μg protein/100 worms/100 μL). The number of live worms was counted daily and plotted on Kaplan Meier curves. Day 0 of the curve corresponds to the day of brain homogenate administration (L4 larval stage of worms). Data are from two independent experiments (30 worms/group). The percentage of live worms seven days after treatment and the median survival were calculated. P >0.05 according to Log-rank and Gehan-Breslow-Wilcoxon tests

